# Does fear of humans predict anti-predator strategies in an ungulate hider species during fawning?

**DOI:** 10.1101/2023.08.16.553188

**Authors:** Jane Faull, Kimberly Conteddu, Laura L. Griffin, Bawan Amin, Adam F. Smith, Amy Haigh, Simone Ciuti

## Abstract

Humans are a major evolutionary force on wildlife via artificial selection. While often explored through the lens of extractive interactions (e.g., hunting) able to favour certain behavioural traits over others, the implications of non-extractive ones, such as wildlife feeding, remain under-studied. Research has recently shown that people tend to feed (and sometimes favour) a limited subset of bolder individuals within natural populations, although its dynamics and consequences are not fully clear. Using fallow deer living in a peri-urban setting as a model population, we studied whether mother deer that display reduced fear of humans and consistently approach them for food adopt weaker anti-predator strategies by selecting for fawning bedsites that are less concealed and closer to human hotspots, allowing them to take advantage of additional artificial feeding opportunities in comparison to shier mothers in this population. Our dataset encompassed 171 fawns from 109 mothers across 4 years. Contrary to our expectations, we found that mothers that regularly accepted food from humans selected for more concealed bedsites farther away from them, giving their offspring better protection while also taking advantage of additional artificial food during lactating. Our results show marked behavioural adaptation by a subset of females, making this the first time that the link between tendency to approach humans and strategies to protect offspring is explored. Given previous findings that these begging females also deliver heavier fawns at birth, our research adds a piece to the complex puzzle describing human manipulation of behaviour in natural populations and its fitness consequences.

## Introduction

As humans and wildlife coexist in all walks of life, we have influenced wildlife behaviour in countless ways over millennia. Due to the ever-increasing human population, associated increases in consumption, and natural environmental cycles, habitats are in a constant state of flux with land use changes like urbanisation and agriculture (McKinney, 2006, Ellis & Ramankutty, 2008). As well as large scale modifications to the landscape, humans can impact wildlife behaviours through their activities and direct interactions with them. Human-wildlife interactions can be classified as extractive, where a resource is removed from the ecosystem by humans, or non-extractive, where interactions occur but nothing is taken away (Griffin and Ciuti, 2023). Extractive interactions like hunting and fishing, where the animal itself is removed from the system, and the impacts of these activities have been extensively researched (Benitez-López et al., 2017, Young et al., 2014).

Wildlife species must now contend with increases in habitat disturbance or loss, scattered or fragmented food sources, changes in ecosystem functioning, frequent human interactions, novel communities and conflict as they move into towns and cities (McKinney, 2002, Sol et al., 2013). These activities have been shown to promote the selection of both behavioural and morphological traits in wildlife (Coltman et al., 2003, Ciuti et al., 2012, Darimont et al., 2009). To successfully live in human dominated landscapes many animals have adopted novel behaviours which allow them to make use of readily available anthropogenically sourced resources (Flemming & Bateman, 2018, Griffin et al. 2022, Sol et al., 2013). It remains largely unknown how behavioural traits can drive the adaptability of wild animals to human-dominated landscapes and how these traits affect a species’ conservation status (Barbosa et al., 2021). Despite these knowledge gaps, there are several examples in the literature of behavioural adaptations wildlife have developed to exploit human environments. Gull species, including Lesser Black-backed Gulls (*Larus fuscus*) and Herring Gulls (*Larus argentatus*), alter their use of foraging patches throughout the day in response to human activity patterns (Spelt et al., 2021). The Australian White Ibis (*Threskiornis molucca*) is known to scavenge through anthropogenic waste, causing management problems locally through pathogen spread (Epstein et al., 2006). Other species have begun to utilise human-derived food sources more directly through artificial feeding activities. In response to climate change and garden feeding, for instance, the Blackcap (*Sylvia atricapilla*) has altered its winter migratory behaviour and now overwinters in Northern Europe (Van Doren et al., 2021).

Artificial feeding has grown in popularity because unlike hunting or fishing, it provides a non-extractive activity that still allows for close interactions with wildlife. It can fall into two categories, accidental and intentional (Milner et al., 2014). Accidental feeding opportunities occur when animals feed on anthropogenic food sources which were not intended for feeding. For example, gulls eating food waste left behind in green areas and parks (Maciusik, Lenda & Skórka, 2010). Additionally, free-ranging dogs (*Canis familiaris*) in less developed countries are known to scavenge on human waste (Butler and du Toit, 2002). Whereas intentional feeding occurs when food is provided with the distinct goal of feeding animals. Most commonly we see examples of people feeding garden birds (Plummer et al., 2019), terrestrial mammals (Knight, 2010, Kojola and Heikkinen, 2012, Usui and Funck, 2018) and marine megafauna (Senigaglia et al., 2020, Fitzpatrick et al., 2011). Human-wildlife feeding research has shown that individuals can be fed as often as every 10 minutes (Usui and Funck, 2018).

While the public perception may be that feeding wildlife is beneficial for animals, as it is non-consumptive and does not involve capturing or killing, research is now focusing on the potentially harmful, unseen impacts (Marion et al., 2008, Dubois and Fraser, 2013, Griffin et al. 2022). The implications of these human activities are bifold as there are consequences for both humans and wildlife (Griffin and Ciuti, 2023). Feeding wildlife has become commonplace, despite being strongly discouraged due to links between wildlife feeding and disease transmission, increased aggression, welfare concerns and personal injury (Marion et al., 2008, Dubois and Fraser, 2013). Many people may naively assume that increased availability of food would lead to improved body condition and overall better health but oftentimes the opposite is true (Mann et al., 2000, Borg et al., 2014, DiMaggio et al., 2023). Considering that the health and fitness of an individual are heavily linked, it is important that we understand how being fed by people impacts an individual’s wellbeing and physiology (Maréchal et al., 2016). Recent research has shown that human presence can elicit a fear response in ungulates similar (or greater) than that seen for natural non-human predators (Crawford et al., 2022) meaning individuals fed in close-contact interactions are likely under increased stress around humans. General parameters of health in Barbary macaques (*Macaca sylvanus*) have shown changes, with human-fed adults displaying larger body sizes and poorer quality coat condition (Borg et al., 2014). Concerningly, the calves of wild dolphins that engage in tourist feeding experience higher mortality than their non-feeding conspecifics (Mann et al., 2000). This illustrates how the behaviour of a parent, its feeding behaviour and anti-predator strategies can impact the health and survival of offspring.

Anti-predator strategies have evolved in wildlife over evolutionary time to allow species to defend and conceal themselves and their offspring against predation threats circumnavigating capture and consumption (Creel et al., 2005). Like any other behaviour, these strategies have become adapted to living in human-dominated landscapes with literature even referring to the “human super predator”, who elicits significant fear responses and altered behaviours in wildlife (Smith et al., 2017, Crawford et al., 2022). As well as functioning as predators within ecosystems, humans can also aid in anti-predation because of the “human shield” effect which allows prey species to live near humans by taking advantage of large predators’ aversion to human presence. This phenomenon has been seen in many predator-prey relationships, including elk (*Cervus canadensis*) and wolves (*Canis lupus*) in the Rocky Mountains, and tapirs (*Tapirus bairdii*) and jaguars (*Panthera onca*) in tropical forests in the Yucatan Peninsula (Hebblewhite et al., 2005, Pérez-Flores et al., 2022). This human shield effect is often utilised when selecting suitable sites to deliver offspring as has been seen in brown bears (*Ursus arctos*) that make use of humans as protective shields for females wishing to defend their cubs from sexually selected infanticide (Steyaert et al., 2016).

We are beginning to understand the impacts on individuals who engage with humans (Griffin et al. 2022, McLaughlin et al. 2022, Griffin et al. 2023), although we do not fully know how these associations can affect future generations via selection pressures over evolutionary time. Looking at the effects of human-wildlife interactions on parental care and offspring fitness can give us an insight into future impacts, should these activities continue, and help to inform management and mitigation strategies. Given the picture we have drawn so far, we investigated anti-predator strategies adopted by females during weaning in an animal population living in a peri-urban area where human-wildlife feeding interactions occur. The Phoenix Park, embedded within the Dublin metropolitan area in Ireland, is Europe’s largest urban park and one such location. Urban and national parklands such as the Phoenix Park provide ample opportunities for hand-feeding and close interaction between visitors and the free-ranging herd of fallow deer (*Dama dama)* (Griffin and Ciuti, 2023). Despite being strongly discouraged by management, feeding the deer has become commonplace in the park (Griffin et al., 2022, Griffin et al., 2023). This popular activity has led researchers to identify a spectrum of distinct behavioural types in the population to date, consisting of bolder “acceptors” and shyer “avoiders” (Griffin et al., 2022). Individuals boldly approaching humans and accepting food (known as consistent beggars) account for ∼20% of the population that tolerate close contact with humans and associate them with food (Griffin et al., 2022). In comparison, shyer individuals have a lower tolerance for human presence, either avoiding engagement with humans attempting to interact or actively moving away from groups of people (Griffin et al. 2022).

Our research specifically aims to disentangle mothers’ anti-predator strategies during the fawning season as a function of their willingness to accept food from humans. In most cases, humans are the main threat for wildlife within human-dominated landscapes (Crawford et al., 2022, Smith et al., 2017). This suggests that anti-predator strategies adopted by fearless mothers, particularly during the birthing season, may be different from those that are more afraid of humans. Our main prediction is that females that are less afraid of humans may have weakened their anti-predator strategies during weaning when living within human dominated areas. Specifically, we predict that bolder mothers that commonly accept food from park visitors would hide their fawns in bedsites closer to human feeding hotspots which would allow them to access feeding opportunities more easily. As feeding occurs within accessible open areas of the park, hiding fawns closer to the hotspots of human feeding may be linked with poorer conditions for fawn concealment. These bedsites may have a higher visibility from most directions, leaving them open to discovery by predators, disturbance and harsh weather conditions. Furthermore, we predict that the link between a mother’s willingness to accept food from humans and bedsite selection (closer to human hotspots) would be stronger during the first days of a fawn’s life – when the decision of where the neonate fawn would be delivered and concealed is expected to be entirely taken by the mother – and weaker for bedsite locations occupied by older, more mobile and independent fawns.

## Materials and methods

### Study site and population

We conducted our study on a free-ranging population of fallow deer inhabiting the largest enclosed urban park of Europe, the 7km^2^ Phoenix Park in Dublin City centre, Ireland (53.3559° N, 6.3298° W) over 4 years (2018-2021). The fallow deer population is maintained at ∼600 individuals by annual culls (targeting ∼75 deer per year, Office of Public Works OPW, official data). Other causes of mortality in this population are traffic collisions and occasional predation upon neonate fawns by foxes (*Vulpes vulpes*) and unleashed dogs. Phoenix Park is a public site with roads and walkways running throughout, making much of the park accessible to the public and providing ample opportunities for human interaction with the deer (Fig. 1, Griffin et al. 2022). The park receives an estimated 10 million visitors annually (OPW, official data). Human-mediated feeding can be opportunistic, whereby visitors offer food that was originally brought for their own consumption (or plants sourced from the park), or premeditated, whereby food items such as carrots have been brought with the intention of feeding the deer (Fig. 1).

**Figure 1:**
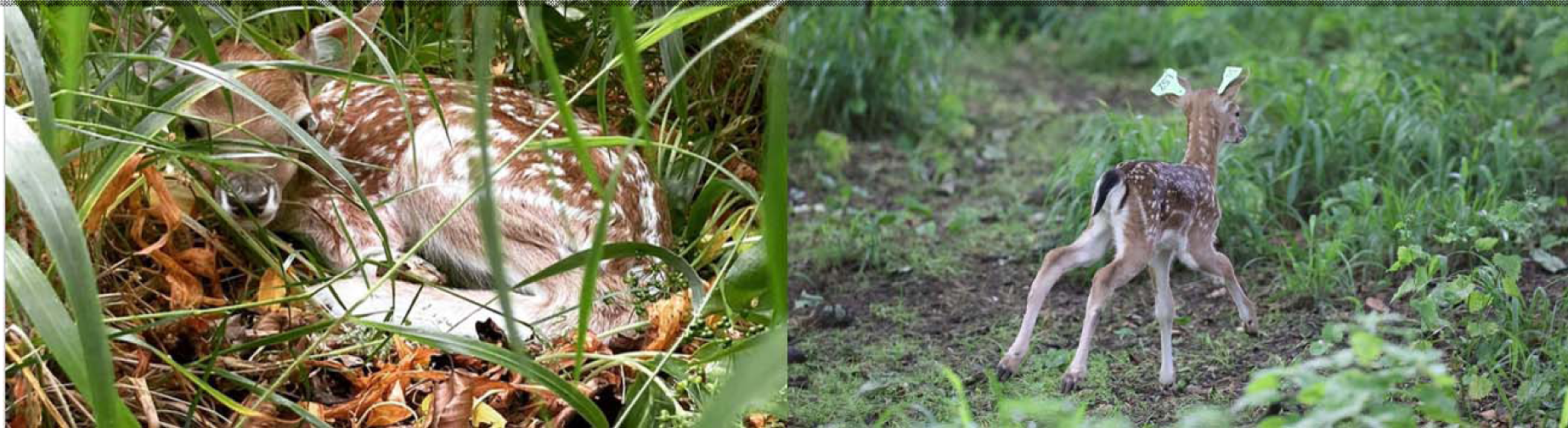
From top left clockwise i) adult females (picture by Laura L. Griffin) and ii) adult males (picture by Bawan Amin) being hand fed carrots by park visitors; iii) neonate fawn hidden in the bedsite vegetation (picture by Clíodhna Hynes) and iv) running away at release after being ear-tagged by University College Dublin capture crew team (picture by Clíodhna Hynes).

### Fawn capture protocol

Fallow deer fawns are ear-tagged (unique colour-number and colour-letter-number combinations) annually during the first three weeks of life, resulting in the vast majority (>80%) of the deer being individually recognisable for management and longitudinal research purposes. Ear-tagging during our study was conducted under animal care permit UCD-AREC-E-18-28, which also covers all non-invasive behavioural observations described in our work. We subdivided the area of our study site that is typically selected by deer mothers to give birth and conceal their fawns into 7 sectors, all of which were patrolled by a trained capture team supervised by a certified wildlife biologist. Fallow deer are a hider species, meaning their fawns are concealed amongst vegetation and left unsupervised in bedsites for long periods of time after birth (Kjellander et al., 2012, Torriani et al., 2006). Depending on the number of bedsites discovered in each area, we visited an average of 2-3 sectors a day, and each sector was patrolled at least once every 3 days. Once a bedsite was discovered, we captured fawns using circular fishing nets (1-1.5 m diameter) with elongated handles (1-1.5 m length). Captured fawns were sexed and ear tagged. The fawn was weighed in a 100L cloth bag using digital scales (resolution: 0.01 kg— Dario Markenartikelvertrieb) before being released. The full capture protocol and description of additional data on neonate fawns can be found in Amin et al. (2021). In most cases fawns were recaptured multiple times (from 1 to 4 times over the fawning period, Amin et al. 2021), meaning that the bedsite locations of fawns of varying ages (from neonate fawns up to 3-week-old fawns) were recorded.

We recorded the bedsite coordinates and determined the average bedsite visibility (*sensu* Bongi et al., 2008 and Amin et al. 2021) using a cardboard square the size of a standing fawn (height: 45cm and width: 36cm) that was sectioned off into multiple equally proportioned triangles. The square was held vertically perpendicular to the ground at the bedsite such that an observer 10m away could record the proportion of the square that was visible (i.e. how many triangles) at a height of 70cm from the ground in four cardinal directions. The height matches that of a red fox (*Vulpes vulpes)*, the only natural predator of these fawns.

### Willingness of female mothers to accept food from humans (a.k.a. begging rank)

Concurrent research in the park gathered observational data on human-deer interactions and ranked deer along a continuum ranging from deer avoiding any interaction with humans to deer consistently begging (i.e. approaching humans) for food (Griffin et al., 2022, Griffin et al., 2023). Data collections on deer begging were carried out from May to July each year (2018-2021) as this is an important time of the annual biological cycle of females (late gestation, birth, early weaning, Ciuti et al., 2006). Females were monitored using a stratified sampling design based on time of the day, day of the week, and area of the park with collections running from dawn to dusk (Griffin et al 2022). The whole park area utilized by deer was subdivided into sectors and patrolled on a strictly scheduled basis. Once a herd was identified, observation team members (at most 3 individuals) identified all deer belonging to the group, collected data on human presence and behaviour (e.g., feeding the deer), and on feeding interactions as they occurred. The start and end time of each interaction, number of people involved, identity of deer involved, what they were fed, how they accepted it (from the hand, thrown to the ground, or from a human’s mouth), and any other deer-human or deer-deer interactions that occurred simultaneously (e.g. petting, harassing, or dominance displays) were recorded. The full procedure for collection of begging interactions is outlined in Griffin et al. (2022). This includes how the individual deer were assigned a begging rank, i.e. the best linear unbiased predictor (BLUPs, Robinson, 1991): this corresponds to the random intercept value of a generalized linear mixed effect model fitted to predict the likelihood of a deer to beg for food (ranging from -2.16 to 5.10) corresponding to the deer willingness to approach humans and accept food after taking into account of group size, time of the day, people present, among the many others confounding factors included (Griffin et al. 2022).

### Mother-fawn pairs

To determine maternal connections, we observed females of this population between July and August after the fawning season. There is no paternal care in this species (Chapman et al., 1997) so fawns will first appear in the female herd with their mothers. As such only maternal relationships can be determined and fathers are unknown. During this time, we recorded interactions such as true suckling, following, and social grooming between mothers and fawns including the ear tags of both individuals involved. True suckling (a.k.a. front suckling) occurs when a fawn feeds from its mother in plain sight of the mother (usually from the front or side) so that she is aware this individual is suckling and lets them. True suckling is distinguished in the field from allosuckling (Roulin, 2002), where a fawn approaches a female that is not its mother from the back and attempts to suckle but is usually driven away by the female once she becomes aware. Following behaviour is distinguishable when a fawn stays very close to a female and almost mirrors her actions by moving in sync or very closely behind her. Social grooming can occur either from mother to fawn or fawn to mother. We confirmed a mother-fawn pair after 3 independent sightings (not occurring in the same observation period) of one or a combination of these interactions (Griffin et al, 2023). We only included fawns that had a confirmed mother-fawn pairing as our aim was to investigate whether a mother’s begging rank affected her bedsite location.

### Data handling and analysis

We combined the individual willingness by deer mothers to beg and accept food from park visitors (begging rank *sensu* Griffin et al. 2022) to data of their respective fawns. Each row of our final dataset corresponded to a unique capture event of a given fawn at a recorded bedsite along with bedsite visibility, distance (in meters) to the most popular hotspot of human feeding in the female sector of the park, identity of the fawn and its weight (in kg) and sex, mother’s identity and age (years old). The age of the mother was exact because all individuals in this population were tagged as neonates. The hotspot of feeding was determined by taking the geometric centroid of the spatial observations of human feeding the deer (*sensu* Griffin et al. 2022), which we used to calculate the distance between such a popular location to the bedsite and test our main hypothesis that begging mothers would conceal their fawns closer to humans. Fawn weight was used as a proxy for age (instead of subjective age estimated in days by fawn handlers) *sensu* Amin et al (2021), being the 2 metrics highly correlated (r_s_ = 0.77).

We used multivariate mixed-effect models to estimate the link between mother’s begging behaviour and the characteristics of the selected bedsites. Using the *brms* package (Buerkner, 2017), we fit a Bayesian multivariate mixed-effect model to explain the variability and covariance of two response variables describing the characteristics of the bedsite (bedsite visibility and its distance to human feeding hotspot) as a function of the following predictors: begging rank and age of the mother; weight of the fawn at capture and its sex. The model was as follows:

> *(Bedsite visibility + Distance to feeding hotspot) ∼ Mother’s begging rank + Fawn weight + Mother’s age + Sex + Mother’s begging rank*Fawn weight + Mother’s age*Fawn weight + Mother ID + Year*

We included the interaction between mothers’ begging rank and fawn weight (proxy for age) to test our hypothesis that lighter (younger) fawns and related bedsite characteristics will be more strongly driven by mothers’ begging rank – as opposed to older fawns which can be more independent, mobile, and less driven by mothers in terms of where to hide. Table 1 provides a summary of all the variables used. We also included the interaction between mothers’ age and fawn weight to control for the fact that more experienced (older) females may conceal their neonate fawns better. All numerical predictors were scaled to improve model convergence. Including both single and quadratic terms of our explanatory variables resulted in increased uncertainty in the model and overfitting. Therefore, based on the inspection of model fit and residuals’ patterns, we were satisfied by the fact that the inclusion of single terms and two-way interactions gave the model sufficient flexibility to account for non-linear effects. Finally, identity of the mother and year of capture were included in the model as crossed random intercepts. All predictors included in the model were successfully screened for collinearity issues (|r_p_| < 0.7) (Dormann et al., 2013). Begging rank of mothers was correlated to their age but far from collinear, with a Pearson correlation coefficient r_p_ = 0.108, showing the tendency of older mother to be more likely to beg for food than younger ones (sensu Griffin et al. 2022). This allowed us to include both predictors in the model and assess the effect of mothers’ begging rank on selected bedsite characteristics while accounting for age and experience. All data handling and analysis, including statistical and GIS analysis was carried out using R 4.0.5. (R Core Team 2021).

**Table 1:** Summary of the variable used in the model. All numerical predictors were scaled to improve model convergence.

| Variable | Description | Scaled |
| --- | --- | --- |
| Bedsite visibility (BV) | Visibility of bedsite to predators | Yes |
| Distance to feeding hotspot | Distance to popular human feeding hotspot within the park | Yes |
| Mother's begging rank | Numerical score measuring how consistently mother accepts food from humans | Yes |
| Fawn Weight | Weight of fawn at capture | Yes |
| Mother's age | Age of mother | Yes |
| Sex | Sex of fawn | No |
| Mother ID | Identity of the mother | No |
| Year | Year of fawn capture | No |

## Results

Data were collected from 280 capture events of 171 neonate fawns born to 109 mothers. The dataset encompasses multiple fawning seasons across 4 subsequent years (2018-2021 inclusive). The n = 171 fawns were evenly distributed across the study with 42 fawns in 2018, 38 in 2019, 44 in 2020 and 47 in 2021. Ninety fawns were captured once, 57 twice, 20 three times, and only 4 four times within the same fawning season. Mother’s age ranged between 2-17 years (mean: 6.7 years) across the study period. Of the 109 mothers included in the study, 63 gave birth to one fawn only, 33 to two, 10 to three and only 3 females gave birth to 4 fawns over 4 years.

We reported in Table 2 the parameters estimated by our Bayesian multivariate mixed-effect model explaining the variability of the two response variables (bedsite visibility and its distance to people feeding hotspot, both fitted with Gaussian distribution of errors) inclusive of the variation of the crossed random intercepts (mother’s identity and year of study). We did not find a clear covariance between bedsite visibility and their distance to the hotspot of people feeding (group-level effects in Table 2), showing that bedsites with low visibility can be found both close and far from the people hotspot. In relation to population-level effects, specifically referring to single effects not included within interactions, we found no effect of the sex of the fawn on the characteristics of the bedsite (both visibility and distance to the human feeding hotspot, Table 2). In relation to the interaction terms specifically included to test our *a priori* hypotheses, we found clear effects of the interactions of mother’s begging ranks with fawn’s weight (proxy for age), as well as of the interaction between mother’s age (proxy for experience) and fawn’s weight (Table 2). We have expanded these results below along with the relevant figures.

**Table 2:** Parameters estimated by the Bayesian multivariate mixed-effect model explaining the covariation of the two response variables (bedsite visibility (BV) and its distance to people feeding hotspot (DH), both fitted with Gaussian distribution of errors) as a function of mother’s begging rank and age, weight and sex of the fawn inclusive of *a priori* interactive effects. Mother identity and year of study were both fitted as crossed random intercepts. The model was fitted on n = 280 observations drawing from 3 chains, each with 4000 iterations (warmup = 250, thin = 2, total post-warmup draws = 5625). To improve readability, asterisks (*) have been added to indicate estimate and related 95% confidence intervals not passing zero.

| **Group-Level Effects** |  |  |  |  |  |  |  |
| --- | --- | --- | --- | --- | --- | --- | --- |
| **~Mother** |  |  |  |  |  |  |  |
|  | **Estimate** | **Est. Error** | **l-95% CI** | **U-95% CI** | **Rh at** | **Bulk ESS** | **Tail ESS** |
| Intercept SD (BV) | 0.21 * | 0.15 | 0.01 | 0.52 | 1 | 1505 | 4993 |
| Intercept SD (DH) | 0.70 * | 0.07 | 0.57 | 0.83 | 1 | 6739 | 9454 |
| Intercept correlation (BV vs DH) | -0.04 | 0.40 | -0.87 | 0.81 | 1 | 418 | 654 |
| **~Year** |  |  |  |  |  |  |  |
| Intercept SD (BV) | 0.52 * | 0.39 | 0.13 | 1.58 | 1 | 5574 | 7641 |
| Intercept SD (DH) | 0.14 * | 0.16 | 0.00 | 0.58 | 1 | 6197 | 6002 |
| Intercept correlation (BV vs DH) | 0.16 | 0.58 | -0.92 | 0.97 | 1 | 9056 | 9826 |
| **Population-Level Effects** |  |  |  |  |  |  |  |
| Intercept (BV) | -0.00 | 0.31 | -0.67 | 0.62 | 1 | 7954 | 7375 |
| Intercept (DH) | 0.02 | 0.14 | -0.25 | 0.29 | 1 | 8588 | 8383 |
| Mother's begging rank (BV) | -0.08 | 0.06 | -0.20 | 0.04 | 1 | 8150 | 9167 |
| Mother's begging rank (DH) | 0.10 | 0.06 | -0.02 | 0.23 | 1 | 9385 | 9474 |
| Fawn's weight (BV) | -0.31 * | 0.06 | -0.43 | -0.19 | 1 | 7627 | 8829 |
| Fawn's weight (DH) | -0.09 * | 0.05 | -0.19 | 0.01 | 1 | 9200 | 9703 |
| Mother's age (BV) | -0.14 * | 0.07 | -0.27 | -0.01 | 1 | 8530 | 9152 |
| Mother's age (DH) | 0.17 * | 0.08 | 0.01 | 0.33 | 1 | 9335 | 9155 |
| Sex of fawn (m) (BV) | -0.05 | 0.12 | -0.29 | 0.18 | 1 | 8287 | 9730 |
| Sex of fawn (m) (DH) | 0.05 | 0.11 | -0.16 | 0.26 | 1 | 9917 | 10060 |
| Mother's begging rank * Fawn's weight (BV) | 0.11 | 0.06 | -0.00 | 0.24 | 1 | 7674 | 9003 |
| Mother's begging rank * Fawn's weight (DH) | -0.13 * | 0.05 | -0.24 | -0.03 | 1 | 8554 | 9795 |
| Fawn's weight * Mother's age (BV) | 0.12 * | 0.05 | 0.02 | 0.23 | 1 | 7994 | 8805 |
| Fawn's weight * Mother's age (DH) | -0.10 * | 0.05 | -0.19 | -0.01 | 1 | 9514 | 9159 |
| **Family specific parameters:** |  |  |  |  |  |  |  |
| Sigma (BV) | 0.90 | 0.05 | 0.80 | 0.99 | 1 | 3306 | 5735 |
| Sigma (DH) | 0.66 | 0.04 | 0.59 | 0.73 | 1 | 7028 | 9016 |

(BV): Bedside visibility is the response variable
(DH): Distance to the human Hotspot is the response variable

We found a strong tendency by consistent beggar mothers (i.e. higher begging rank) to conceal their fawns in sites with reduced visibility (Table 2; Fig. 2, left panel), but this was true for younger (lighter) fawns and not for older (heavier) and more mobile fawns. Contrary to our main expectations, fawns of consistent beggar mothers were found in areas further away from the hotspot of human feeding when compared to the fawns of shyer mothers with lower begging ranks (Table 2; Fig 3, left panel). This pattern was, again, clear when the fawns were younger (lighter) and vanished for older and heavier fawns (Fig. 3 left panel).

**Figure 2:**
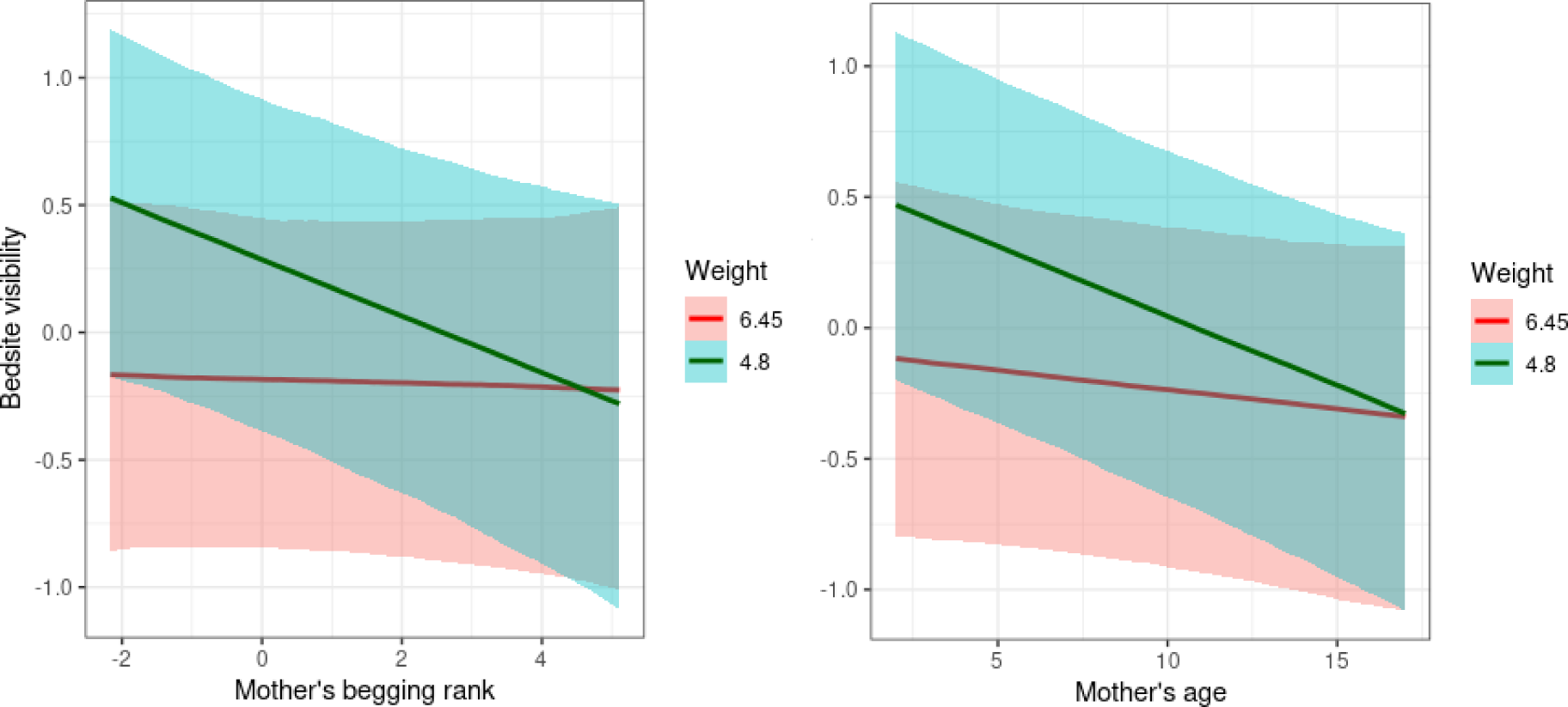
Effect of begging rank of the mother (x-axis, left panel) and mother’s age (years old, x-axis, right panel) interacted with the weight of the fawn (first and third quartile, in kg) on the visibility (scaled, y-axis) of bedsites as predicted by the Bayesian multivariate mixed-effect model. Begging mothers (higher begging rank) as well as older and more experienced mothers tended to hide their fawns in less visible sites (lower visibility) when the fawns were younger (lighter weight), whereas such a relationship was absent for heavier (older) fawns.

**Figure 3:**
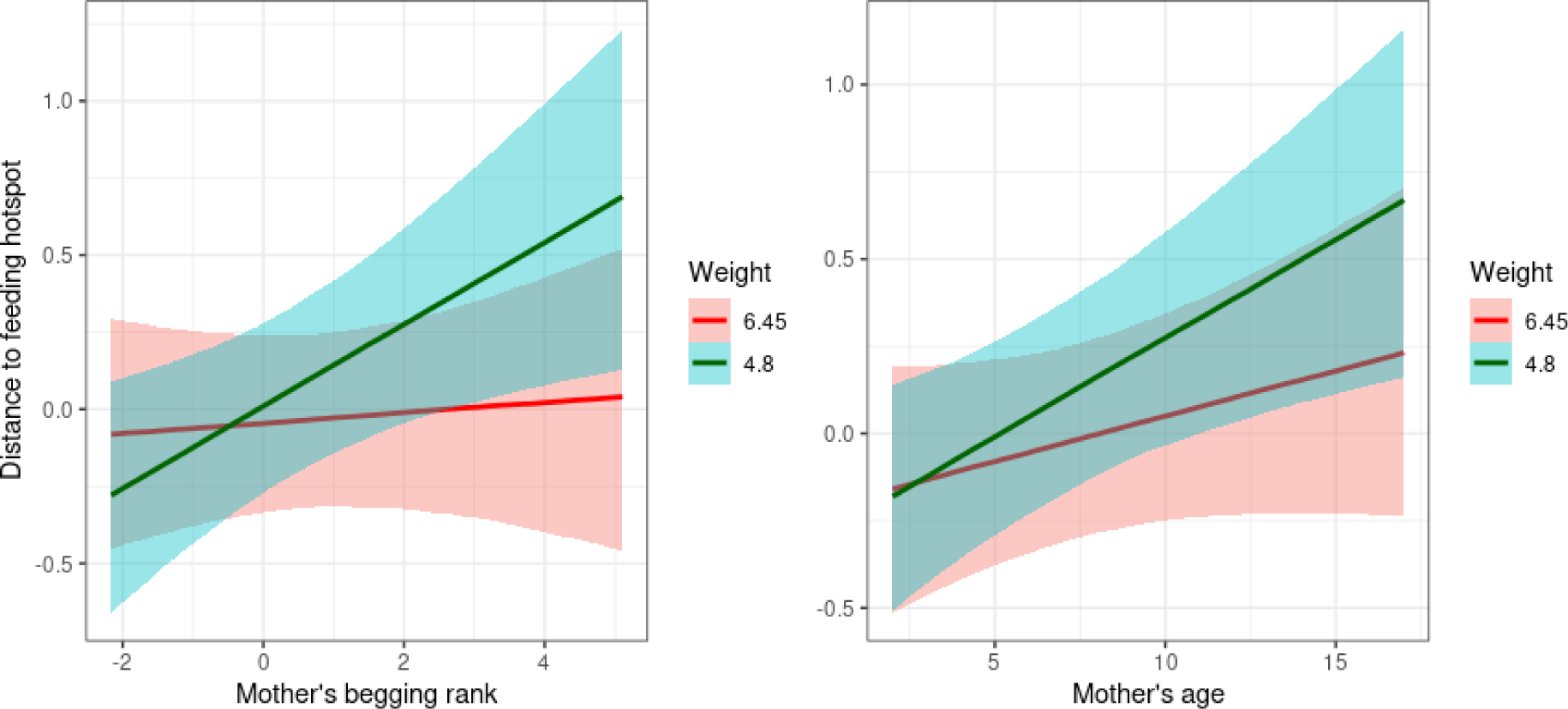
Effect of begging rank of the mother (x-axis, left panel) and mother’s age (years old, x-axis, right panel) interacted with the weight of the fawn (first and third quartile, in kg) on the distance to feeding hotspot (scaled, y-axis) of bedsites as predicted by the Bayesian multivariate mixed-effect model. Fawns were hidden in sites further from feeding hotspots by both consistent begging mothers and older and more experienced ones, whereas such a relationship was absent for heavier (older) fawns. Note that in the plot a scaled value of distance 1 corresponds to 901m and a scaled distance of -0.5 corresponds to 476m.

When looking at the interaction of mother’s age and fawn weights (Table 2), accounting for the effect of begging rank, older and more experienced mothers concealed their fawns in less visible sites (Fig. 2 right panel) and more distant to the hotspot of human feeding (Fig. 3 right panel). Similarly to the interaction with mother’s begging rank, the pattern was evident in younger (lighter) fawns and vanished in older and heavier fawns.

## Discussion

Our aim was to disentangle anti-predator strategies during the birth period of consistently begging mothers’ in a semi-urban environment. Our main *a priori* expectation was that female deer that show a reduced fear response to humans and regularly accept food from them (sensu Griffin et al. 2022) would have concealed their fawns closer to the hotspot of artificial feeding. By doing so they could more easily exploit artificial food while remaining relatively close to their concealed offspring, facilitating maternal care responsibilities such as suckling. Contrary to our expectation, consistent beggar females displayed the most adaptable behaviour within a relatively small site with high human activity. Consistent beggar mothers took advantage of artificial feeding opportunities by showing higher acceptance rates than others in the population (Griffin et al. 2022) and as we have shown in our study, they concealed their fawns in areas that tended to have thicker vegetation and were farther away from human feeding hotspots compared to shyer females. Our results clearly document a remarkable adaptation to local ecological conditions shown by a subset of female deer living in an urban park. These findings raise questions pertaining to how inter-individual differences in response to human selection pressures may shape the behaviour of wildlife within increasingly human-dominated landscapes. Our work adds an important piece to this puzzle - describing in detail inter-individual variability in coping with humans within the same population, but also raises new questions which we expand upon further below.

The same preferences for birthing and hiding fawns in concealed areas with thicker vegetation further away from human feeding hotspots shown by begging females are adopted by older mothers. In general, older females will have gained experience from raising previous offspring and, therefore, females tend to become better mothers as they age (Paitz et al., 2007). But consistent beggar mothers vary across the age range within our population with some being much younger and less experienced mothers than others. This illustrates that individuals engaging in begging activities have acquired beneficial behavioural traits normally associated with older, wiser, and more experienced mothers.

We *a priori* predicted that the link between mothers’ willingness to accept food from humans and bedsite selection closer to human hotspots would be stronger during the first days of a fawn’s life – when the decision of where the neonate fawn will be delivered and concealed is expected to be entirely taken by the mother – and weaker for bedsite locations occupied by older, more mobile, and independent fawns. Our data confirmed this prediction, and we showed that the link between bedsite characteristics and mothers’ behaviour were evident only in captures of young neonates usually within a week after birth, meaning that the decision to choose this location was driven by the mother (Blank, 2017). As fawns mature, mothers begin using contact calls to locate them, suggesting that they have only an approximate knowledge of the fawn’s location (Torriani et al., 2006). This suggests that fawns begin to take an active role in deciding their location and move around more independently of their mothers after the first period of the hiding phase when they barely move.

Bedsite selection is an incredibly important behavioural decision by mothers of hider species because fawn survival would depend on multiple factors including predation, disease, and hypothermia (Jarnemo and Liberg, 2005, Kjellander et al., 2012). The mothers can limit these threats by selecting suitable bedsite locations especially for hider species where a fawn’s greatest defence from predation is cover and cryptic colouration (Grovenburg et al., 2010), and adequate shelter from cold and damp weather conditions can help to fend off hypothermia and disease. It has previously been shown that bolder mothers (the same females included in this study) receive more food than their shy conspecifics, resulting in them giving birth to fawns that are typically 300-500g heavier (Griffin et al., 2022, Griffin et al., 2023). Not only are these fawns heavier, but they also exhibit higher growth rates than the offspring of mothers who do not beg (Griffin et al., 2023). This provides their offspring with an advantage early in life when weight is an important predictor for neonate survival (Amin et al., 2022) and mortality is at its highest in early life for ungulate species (Gaillard et al., 1997). Considering neonate fawns rely solely on their mother’s milk for nutrition in early life (Chapman et al., 1997), this could be linked to previous research which has shown that lactating females who receive supplementary feeding show increased milk production (Garg et al., 2013). Previous research has also shown that prolonged nursing occurs in heavier offspring in similar ungulates (Martinez et al., 2009). Combining these previously established early life characteristics of higher birth weight and faster growth rates (Amin et al., 2022, Griffin et al., 2022, Griffin et al., 2023) with our findings about superior bedsite location, the offspring of bolder mothers are consistently awarded multiple survival advantages from in utero to young adult life stages. This provides evidence that begging, risk taking mothers are better adapted to this environment than shyer, risk avoiding females.

While individuals who regularly engage in human feeding activities may be better adapted in this context, it is important to note that bolder behaviour is not always advantageous as previously mentioned (Mann et al., 2000). The link between increased offspring survival and food acceptance is context dependent. It has been argued that feeding activities should only be deemed acceptable, “if it could be controlled, if it has a beneficial conservation effect and if it did not compromise an animal’s long-term welfare” (Dubois and Fraser, 2013). Another issue with artificial feeding in this scenario is that deer are often given foods that are vastly different from their natural diets (e.g. chocolate, crisps/chips, and sandwiches). When an animal is regularly exposed to human food that they cannot digest it worsens their physical health leaving them more susceptible to parasites and disease (Brookhouse et al., 2013) that could be passed on to other healthy (potentially non-begging) members of the population. Adaptability of species to urban settings has been increasingly studied over the last decade (e.g. Bateman and Fleming 2012; Odden et al. 2014), and concerns have been raised about the non-random sorting of individuals (Sol et al 2013) with the urgency to improve our understanding of the mechanisms through which behaviour helps animals to cope with such environmental alterations.

While bold behaviour may be rewarded in some cases within an intraspecies context through fitness advantages over shyer conspecifics (Amin et al., 2022, Griffin et al., 2022, Griffin et al., 2023), it is questionable whether it is beneficial when we consider inter-species relations. In this landscape, humans and deer are constantly in close proximity due to the size and use of our study site. Bold behavioural types have been linked to behaviours like aggression and risk-taking (Marion et al., 2008, Dubois and Fraser, 2013) which can cause injury (both to deer and people). These types of interactions can lead to increased conflict between humans and wildlife, which in turn may require more robust management strategies (and financial investment) to monitor and minimise these problems.

Previous studies have shown that extractive activities like hunting and fishing are pushing selection towards less desirable traits in wild animals (Allendorf and Hard, 2009). Based on our bedsite selection findings coupled with previous findings of increased fawn birth weights and growth rates (Amin et al.,2022, Griffin et al., 2022, Griffin et al., 2023), beggar individuals have a clear survival advantage over conspecifics that beg less. It could be argued feeding activities promote the artificial selection of bolder begging behaviours (Griffin et al., 2022, Griffin and Ciuti, 2023). In a more natural setting, this behavioural type would exist within a herd but the proportion of bold to shy individuals would be maintained through the associated costs of boldness e.g. predation (Hulthén et al., 2017). However due to the lack of predation for bolder adults in this circumstance boldness can continually be rewarded with additional food without the same level of associated risk which could therefore be encouraging this behaviour. If artificial selection is occurring, and bolder mother’s offspring are surviving better than their shyer conspecifics we could see an increase in the proportion of bold individuals in populations over time.

Our research advances our understanding of human-wildlife interactions and related knock-on effects, but several questions remain unanswered. The next steps in better understanding the impacts of these close contact associations between humans and wildlife are to allocate research efforts into understanding whether these behaviours are passed down through generations and the mechanisms involved. Griffin et al (2022) documented high intra-individual repeatability in fallow deer begging behaviour across years, suggesting that personality could play a role in driving the behaviour of the bold beggars. This would be particularly problematic if the innate propensity to interact with humans is a heritable trait like other personality dimensions (e.g. Dochtermann et al. 2015, Sinn et al. 2006, Weiss et al. 2000). It is not yet understood whether this propensity would be inherited genetically or – alternatively or in addition - if cultural transmission of begging behaviours occurs from mother to offspring, either or both contributing to the maintenance and potential increase in the frequency of this behaviour over generations. Either way, it must be a research priority to better understand how humans are – voluntarily or not - shaping the behaviour of wildlife within increasingly human dominated landscapes.

## Acknowledgements

We wish to express our sincerest gratitude to all the field workers and volunteers who helped in collection of data during fawn capture and behavioural observations. We thank the Office of Public Works, Ireland, for support (grant R19730). We also thank the School of Biology and Environmental Sciences in University College Dublin and the HEA for the associated co-funding. This publication has emanated from research conducted with the financial support of Science Foundation Ireland under Grant number 18/CRT/6049. For the purpose of Open Access, the author has applied a CC BY public copyright licence to any Author Accepted Manuscript version arising from this submission.

## Notes

### Competing Interest Statement

The authors have declared no competing interest.

## Bibliography

Allendorf, F. W. & Hard, J. J. 2009. Human-induced evolution caused by unnatural selection through harvest of wild animals. Proceedings of the National Academy of Sciences - PNAS, vol. 106, pp. 9987–9994.

Amin, B., Jennings, D.J., Smith, A.F., Quinn, M., Chari, S., Haigh, A., Matas, D., Koren, L. & Ciuti, S. 2021, “In utero accumulated steroids predict neonate antilJpredator response in a wild mammal”, Functional ecology, vol. 35, no. 6, pp. 1255–1267.

Amin, B., Verbeek, L., Haigh, A., Griffin, L.L. & Ciuti, S. 2022, “Risk-taking neonates do not pay a survival cost in a free-ranging large mammal, the fallow deer (*Dama dama*)”, Royal Society open science, vol. 9, no. 9, pp. 220578–220578

Barbosa, J.M., Hiraldo, F., Romero, M.Á. & Tella, J.L. 2021, “When does agriculture enter into conflict with wildlife? A global assessment of parrot–agriculture conflicts and their conservation effects”, Diversity & distributions, vol. 27, no. 1, pp. 4–17.

Bateman, P.W. & Fleming, P.A. 2012, “Big city life: carnivores in urban environments”, Journal of zoology (1987), vol. 287, no. 1, pp. 1-23.

Benítez-López, A., Alkemade, R., Schipper, A.M., Ingram, D.J., Verweij, P.A., Eikelboom, J.A.J. & Huijbregts, M.A.J. 2017, “The impact of hunting on tropical mammal and bird populations”, Science (American Association for the Advancement of Science), vol. 356, no. 6334, pp. 180–183.

Blank, D. 2017, “Antipredator tactics are largely maternally controlled in goitered gazelle, a hider ungulate”, Behavioural processes, vol. 136, pp. 28–35.

Bongi, P., Ciuti, S., Grignolio, S., Del Frate, M., Simi, S., Gandelli, D. & Apollonio, M. 2008, “Anti-predator behaviour, space use and habitat selection in female roe deer during the fawning season in a wolf area”, Journal of zoology (1987), vol. 276, no. 3, pp. 242-251.

Borg, C., Majolo, B., Qarro, M. & Semple, S. 2014. “A Comparison of Body Size, Coat Condition and Endoparasite Diversity of Wild Barbary Macaques Exposed to Different Levels of Tourism”, Anthrozoös, vol. 27, pp. 49–63.

Brookhouse, N., Buchera, D. J., Roseb, K., Kerrc, I. & Gudge, S. 2013. Impacts, risks and management of fish feeding at Neds Beach, Lord Howe Island Marine Park, Australia: a case study of how a seemingly innocuous activity can become a serious problem. Journal of Ecotourism, Vol. 12, pp. 165–181.

Buerkner, P. C. 2017. brms: An R package for Bayesian multilevel models using Stan. Journal of statistical software, vol. 80, pp. 1–28.

Butler, J.R.A. & Du Toit, J.T. 2002, “Diet of free-ranging domestic dogs (*Canis familiaris*) in rural Zimbabwe: implications for wild scavengers on the periphery of wildlife reserves”, Animal conservation, vol. 5, no. 1, pp. 29–37.

Chapman, D., Chapman, N. & Fawcett, J. K. 1997. Fallow deer: their history, distribution and biology, 2^nd^ edn, Lavenham, Terence Dalton Limited.

Ciuti, S., Bongi, P., Vassale, S. & Apollonio, M. 2006. Influence of fawning on the spatial behaviour and habitat selection of female fallow deer (*Dama dama*) during late pregnancy and early lactation. Journal of Zoology, 268, 97–107.

Ciuti, S., Muhly, T.B., Paton, D.G., Mcdevitt, A.D., Musiani, M. & Boyce, M.S. 2012; 1107; “Human selection of elk behavioural traits in a landscape of fear”, *Proceedings of the Royal Society. B*, Biological sciences, vol. 279, no. 1746, pp. 4407–4416.

Coltman, D.W., O’donoghue, P., Jorgenson, J.T., Hogg, J.T., Strobeck, C. & Festa-Bianchet, M. 2003, “Undesirable evolutionary consequences of trophy hunting”, Nature, vol. 426, no. 6967, pp. 655–658.

Couch, C. E., Wise, B. L., Scurlock, B. M., Rogerson, J. D., Fuda, R. K., Cole, E. K., Szcodronski, K. E., Sepulveda, A. J., Hutchins, P. R. & Cross, P. C. 2021. Effects of supplemental feeding on the fecal bacterial communities of Rocky Mountain elk in the Greater Yellowstone Ecosystem. PloS one, vol. 16, no. 4, e0249521–e0249521.

Crawford, D. A., Conner, L. M., Clinchy, M., Zanette, L. Y. & Cherry, M. J. 2022. Prey tells, large herbivores fear the human ‘super predator’. Oecologia, vol. 198, pp. 91–98.

Creel, S., Winnie, J., Maxwell, B., Hamlin, K. & Creel, M. 2005, “Elk alter habitat selection as an antipredator response to wolves”, Ecology (Durham), vol. 86, no. 12, pp. 3387–3397.

Darimont, C.T., Carlson, S.M., Kinnison, M.T., Paquet, P.C., Reimchen, T.E., Wilmers, C.C. & Daily, G.C. 2009, “Human Predators Outpace Other Agents of Trait Change in the Wild”, Proceedings of the National Academy of Sciences - PNAS, vol. 106, no. 3, pp. 952–954.

Dimaggio, K.M., Acevedo, M.A., Mchugh, K.A., Wilkinson, K.A., Allen, J.B. & Wells, R.S. 2023, “The fitness consequences of humanlJwildlife interactions on foraging common bottlenose dolphins (*Tursiops truncatus*) in Sarasota Bay, Florida”, Marine mammal science, 1-17.

Dochtermann, N.A., Schwab, T. & Sih, A. 2015, “The contribution of additive genetic variation to personality variation: heritability of personality”, *Proceedings of the Royal Society. B*, Biological sciences, vol. 282, no. 1798, pp. 20142201–20142201.

Donaldson, R., Finn, H., Bejder, L., Lusseau, D., Calver, M., Gompper, M. & Williams, R. 2012. The social side of human-wildlife interaction: wildlife can learn harmful behaviours from each other. Animal Conservation, 15, 427–435.

Dormann, C. F., Elith, J., Bacher, S., Buchmann, C., Carl, G., Carré, G., Marquéz, J. R. G., Gruber, B., Lafourcade, B., Leitão, P. J., Münkemüller, T., Mcclean, C., Osborne, P. E., Reineking, B., Schröder, B., Skidmore, A. K., Zurell, D. & Lautenbach, S. 2013. Collinearity: a review of methods to deal with it and a simulation study evaluating their performance. Ecography (Copenhagen), vol. 36, pp. 27–46.

Dubois, S. & Fraser, D. 2013. A framework to evaluate wildlife feeding in research, wildlife management, tourism and recreation. Animals (Basel), vol. 3, pp. 978–994.

Ellis, E.C. & Ramankutty, N. 2008, “Putting people in the map: anthropogenic biomes of the world”, Frontiers in ecology and the environment, vol. 6, no. 8, pp. 439–447.

Epstein, J. H., Mckee, J., Shaw, P., Hicks, V., Micalizzi, G., Daszak, P., Kilpatrick, A. M. & Kaufman, G. 2006. Australian White Ibis (*Threskiornis molucca*) as a Reservoir of Zoonotic and Livestock Pathogens. EcoHealth, vol. 3, pp. 290–298.

Fitzpatrick, R., Abrantes, K. G., Seymour, J. & Barnett, A. 2011. Variation in depth of whitetip reef sharks: does provisioning ecotourism change their behaviour? Coral reefs, vol. 30, pp. 569–577.

Fleming, P.A. & Bateman, P.W. 2018. “Novel predation opportunities in anthropogenic landscapes”, Animal behaviour, vol. 138, pp. 145–155.

Gaillard, J., Boutin, J., Delorme, D., Van Laere, G., Duncan, P. & Lebreton, J. 1997, “Early Survival in Roe Deer: Causes and Consequences of Cohort Variation in Two Contrasted Populations”, Oecologia, vol. 112, no. 4, pp. 502–513.

Garg, M.R., Sherasia, P.L., Bhanderi, B.M., Phondba, B.T., Shelke, S.K. & Makkar, H.P.S. 2013, “Effects of feeding nutritionally balanced rations on animal productivity, feed conversion efficiency, feed nitrogen use efficiency, rumen microbial protein supply, parasitic load, immunity and enteric methane emissions of milking animals under field conditions”, Animal feed science and technology, vol. 179, no. 1-4, pp. 24–35.

Griffin, L.L. & Ciuti, S. 2023, “Should we feed wildlife? A call for further research into this recreational activity”, Conservation science and practice, vol. 5, no. 7, pp. n/a.

Griffin, L. L., Haigh, A., Amin, B., Faull, J., Norman, A. & Ciuti, S. 2022. Artificial selection in humanlJwildlife feeding interactions. Journal of Animal Ecology.

Griffin, Laura L., Haigh, A., Amin, B., Faull, J., Corcoran, F., Baker-Horne, C. & Ciuti, S. 2023, “Does artificial feeding impact neonate growth rates in a large free-ranging mammal?”, Royal Society open science, vol. 10, no. 3.

Grovenburg, T.W., Jacques, C.N., Klaver, R.W. & Jenks, J.A. 2010, “Bed Site Selection by Neonate Deer in Grassland Habitats on the Northern Great Plains”, The Journal of wildlife management, vol. 74, no. 6, pp. 1250–1256.

Hebblewhite, M., Merrill, E.H. & Mcdonald, T.L. 2005, “Spatial decomposition of predation risk using resource selection functions: an example in a wolf-elk predator-prey system”, Oikos, vol. 111, no. 1, pp. 101–111.

Hulthén, K., Chapman, B.B., Nilsson, P.A., Hansson, L., Skov, C., Brodersen, J., Vinterstare, J. & Brönmark, C. 2017, “A predation cost to bold fish in the wild”, Scientific reports, vol. 7, no. 1, pp. 1239–5.

Jarnemo, A. & Liberg, O. 2005, “RED FOX REMOVAL AND ROE DEER FAWN SURVIVAL—A 14-YEAR STUDY”, The Journal of wildlife management, vol. 69, no. 3, pp. 1090–1098.

Kjellander, P., Svartholm, I., Bergvall, U.A. & Jarnemo, A. 2012, “Habitat use, bed-site selection and mortality rate in neonate fallow deer *Dama dama*”, Wildlife biology, vol. 18, no. 3, pp. 280–291.

Knight, J. 2010. The ready-to-view wild monkey. The convenience principle in Japanese wildlife tourism. Annals of tourism research, vol. 37, pp. 744-762.

Kojola, I. & Heikkinen, S. 2012. Problem brown bears Ursus arctos in Finland in relation to bear feeding for tourism purposes and the density of bears and humans. Wildlife biology, vol. 18, pp. 258–263.

Maciusik, B., Lenda, M. & Skórka, P. 2010, “Corridors, local food resources, and climatic conditions affect the utilization of the urban environment by the Black-headed Gull *Larus ridibundus* in winter”, Ecological research, vol. 25, no. 2, pp. 263–272.

Mann, J., Connor, R. C., Barre, L. M. & Heithaus, M. R. 2000. Female reproductive success in bottlenose dolphins (*Tursiops sp*.): life history, habitat, provisioning, and group-size effects. Behavioral ecology, vol. 11, pp. 210–219.

Maréchal, L., Semple, S., Majolo, B. & Maclarnon, A. 2016. Assessing the effects of tourist provisioning on the health of wild Barbary macaques in Morocco. PloS one, vol. 11, pp. e0155920–e0155920.

Marion, J. L., Dvorak, R. G. & Manning, R. E. 2008. Wildlife Feeding in Parks: Methods for Monitoring the Effectiveness of Educational Interventions and Wildlife Food Attraction Behaviors., Human Dimensions of Wildlife, vol. 13, pp. 429–442.

Martínez, M., Otal, J., Ramírez, A., Hevia, M.L. & Quiles, A. 2009, “Variability in the behavior of kids born of primiparous goats during the first hour after parturition: Effect of the type of parturition, sex, duration of birth, and maternal behavior”, Journal of animal science, vol. 87, no. 5, pp. 1772–1777.

Mclaughlin, D., Griffin, L.L., Ciuti, S. & Stewart, G. 2022, “Wildlife feeding activities induce papillae proliferation in the rumen of fallow deer”, Mammal research, vol. 67, no. 4, pp. 525–530.

Mckinney, M. L. 2002. “Urbanization, Biodiversity, and Conservation”, Bioscience, vol. 52, pp. 883–890.

Mckinney, M. L. 2006. “Urbanization as a major cause of biotic homogenization”, Biological conservation, vol. 127, pp. 247–260.

Milner, J.M., Van Beest, F.M., Schmidt, K.T., Brook, R.K. & Storaas, T. 2014, “To feed or not to feed? Evidence of the intended and unintended effects of feeding wild ungulates”, The Journal of wildlife management, vol. 78, no. 8, pp. 1322–1334.

Odden, M., Athreya, V., Rattan, S. & Linnell, J.D.C. 2014, “Adaptable neighbours: Movement patterns of GPS-collared leopards in human dominated landscapes in India”, PloS one, vol. 9, no. 11, pp. e112044–e112044.

Paitz, R.T., Harms, H.K., Bowden, R.M. & Janzen, F.J. 2007, “Experience pays: offspring survival increases with female age”, Biology letters (2005), vol. 3, no. 1, pp. 44–46.

Pérez-Flores, J., Hénaut, Y., Sanvicente, M., Pablo-Rodríguez, N. & Calmé, S. 2022, “Jaguar’s Predation and Human Shield, a Tapir Story”, Diversity, vol. 14, no. 12, pp. 1103.

Plummer, K. E., Risely, K., Toms, M. P. & Siriwardena, G. M. 2019. The composition of British bird communities is associated with long-term garden bird feeding. Nature communications, vol. 10, pp. 2088–2088.

R CORE TEAM 2021. R: A language and environment for statistical computing. R Foundation for Statistical Computing, Vienna, Austria

Robinson, G. K. 1991. That BLUP is a good thing: the estimation of random effects. Statistical science, pp. 15–32.

Roulin, A. 2002, Why do lactating females nurse alien offspring? A review of hypotheses and empirical evidence, Elsevier Ltd, London.

Senigaglia, V., New, L. & Hughes, M. 2020. Close encounters of the dolphin kind: Contrasting tourist support for feeding based interactions with concern for dolphin welfare. Tourism management (1982), vol. 77, pp. 104007.

Sinn, D.L., Apiolaza, L.A. & Moltschaniwskyj, N.A. 2006, “Heritability and fitnesslJrelated consequences of squid personality traits”, Journal of evolutionary biology, vol. 19, no. 5, pp. 1437–1447.

Smith, J.A., Suraci, J.P., Clinchy, M., Crawford, A., Roberts, D., Zanette, L.Y. & Wilmers, C.C. 2017, “Fear of the human ‘super predator’ reduces feeding time in large carnivores”, *Proceedings of the Royal Society. B*, Biological sciences, vol. 284, no. 1857, pp. 20170433–20170433.

Sol, D., Lapiedra, O. & González-Lagos, C. 2013. Behavioural adjustments for a life in the city. Animal behaviour, vol. 85, pp. 1101–1112.

Spelt, A., Soutar, O., Williamson, C., Memmott, J., Shamoun-Baranes, J., Rock, P. & Windsor, S. 2021. Urban gulls adapt foraging schedule to human-activity patterns. *Ibis (London*, England), vol. 163, pp. 274–282.

Steyaert, S. M. J. G., Leclerc, M., Pelletier, F., Kindberg, J., Brunberg, S., Swenson, J.E., Zedrosser, A. & Sveriges Lantbruksuniversitet 2016, “Human shields mediate sexual conflict in a top predator”, *Proceedings of the Royal Society. B*, Biological sciences, vol. 283, no. 1833, pp. 20160906–20160906.

Torriani, M., Vannoni, E. & Mcelligott, A. 2006, “MotherlJYoung Recognition in an Ungulate Hider Species: A Unidirectional Process”, The American naturalist, vol. 168, no. 3, pp. 412–420.

Usui, R. & Funck, C. 2018. Analysing food-derived interactions between tourists and sika deer (*Cervus nippon*) at Miyajima Island in Hiroshima, Japan: implications for the physical health of deer in an anthropogenic environment. Journal of Ecotourism, Vol. 17, pp. 67–78.

Van Doren, B.M., Conway, G.J., Phillips, R.J., Evans, G.C., Roberts, G.C.M., Liedvogel, M. & Sheldon, B.C. 2021, “Human activity shapes the wintering ecology of a migratory bird”, Global change biology, vol. 27, no. 12, pp. 2715–2727.

Weiss, A., King, J.E. & Figueredo, A.J. 2000, “The heritability of personality factors in chimpanzees (Pan troglodytes)”, Behavior genetics, vol. 30, no. 3, pp. 213–221.

Young, M.A.L., Foale, S. & Bellwood, D.R. 2014, “Impacts of recreational fishing in Australia: historical declines, self-regulation and evidence of an early warning system”, Environmental conservation, vol. 41, no. 4, pp. 350–356.

